# True UV color vision in a butterfly with two UV opsins

**DOI:** 10.1101/2020.11.14.382507

**Authors:** Susan D. Finkbeiner, Adriana D. Briscoe

## Abstract

1. True color vision in animals is achieved when wavelength discrimination occurs based on chromatic content of the stimuli, regardless of intensity. In order to successfully discriminate between multiple wavelengths, animals must use at least two photoreceptor types with different spectral sensitivity peaks.
2. *Heliconius* butterflies have duplicate UV opsin genes, which encode two kinds of photoreceptors with peak sensitivities in the ultraviolet and violet, respectively. In *H. erato*, the ultraviolet photoreceptor is only expressed in females.
3. Evidence from intracellular recordings suggests female *H. erato* may be able to discriminate between UV wavelengths, however, this has yet to be tested experimentally.
4. Using an arena with a controlled light setting, we tested the ability of *H. erato*, and two species lacking the violet receptor, *H. melpomene* and outgroup *Eueides isabella*, to discriminate between two ultraviolet wavelengths, 380 and 390 nm, as well as two blue wavelengths, 400 and 436 nm, after being trained to associate each stimulus with a food reward. Wavelength stimuli were presented in varying intensities to rule out brightness as a cue.
5. We found that *H. erato* females were the only butterflies capable of color vision in the UV range; the other butterflies had an intensity-dependent preference for UV stimuli. Across species, both sexes showed color vision in the blue-range.
6. Models of *H. erato* color vision suggest that females have an advantage over males in discriminating the inner UV-yellow corolla of *Psiguria* pollen flowers from the surrounding outer orange petals, while previous models (McCulloch et al. 2017) suggested that *H. erato* males have an advantage over females in discriminating *Heliconius* 3-hyroxykynurenine (3-OHK) yellow wing coloration from non-3-OHK yellow wing coloration found in mimics.
7. These results provide some of the first behavioral evidence for UV color discrimination in *Heliconius* females in the context of foraging, lending support to the hypothesis (Briscoe et al. 2010) that the duplicated UV opsin genes function together in UV color vision. Taken together, the sexually dimorphic visual system of *H. erato* appears to have been shaped by both sexual selection and sex-specific natural selection.

## INTRODUCTION

Color vision in animals is characterized by wavelength discrimination based on spectral composition of the stimuli, independent of intensity (Kelber and Pfaff 1999). Animals that have true color vision must use at least two types of photoreceptor, with different spectral sensitivities, to successfully discriminate between wavelengths where their sensitivity curves overlap.

Insects use color vision for multiple tasks including foraging (Spaethe et al. 2001, Muth et al. 2015), host plant detection (Scherer and Kolb 1987), and conspecific recognition (Kemp and Rutowski 2011). Most insects have at least one ultraviolet, one blue, and one green photoreceptor, many insects lack red receptors (Briscoe and Chittka 2001), and some have lost their blue receptors (Sharkey et al. 2017). Numerous butterflies, however, have visual systems with more than three photoreceptor classes (van der Kooi et al. 2021).

While butterflies typically have only one kind of UV opsin (Briscoe et al. 2003, Koshitaka et al. 2008), and variable numbers of blue and green opsins, *Heliconius* have duplicated UV opsins (Briscoe et al. 2010). The two UV opsin-encoded photoreceptors have peak sensitivities or λ_max_ values at 355 and 390 nm as measured by intracellular recordings (McCulloch et al. 2016a). Although the gene encoding UVRh2, which together with the chromophore produces a violet receptor, is present throughout the genus, the UVRh2 protein, is only expressed at detectable levels in the eye in certain *Heliconius* clades (specifically the *sara* and *erato* clades); it is also sex-specific. In *H. erato*, adult females express both UVRh1 and UVRh2 opsins but males only express the violet receptor with sensitivity at 390 nm (McCulloch et al. 2017).

*Heliconius* butterflies also have a genus-specific wing pigment, 3-hydroxy-DL-kynurenine (3-OHK), found in the yellow scales of the wings (Brown 1967). Together, the pigment and the wing ultrastructure reflect UV light in the 300-400 nm range and have a step-like reflectance starting about 440 nm. This wing pigment has evolved in *Heliconius* along with their duplicated UV opsins (Briscoe et al. 2010), and close relatives to this genus lack both the opsin duplication and the 3-OHK wing pigment (Yuan et al. 2010). It has been proposed that the second UV opsin might allow for better discrimination of yellow-winged *Heliconius* conspecifics from yellow-winged non-*Heliconius* mimics (Bybee et al. 2012), perhaps because these butterflies’ yellows differ with respect to UV wavelengths; recent experiments lend some support to this hypothesis (Finkbeiner et al. 2017, Dell’Aglio et al. 2018) but more behavioral experiments examining the functional significance of the duplicate UV opsins are needed.

In *Heliconius* or passion-vine butterflies, adults have large heads relative to body mass (compared to other butterflies) with notable investment in the visual neuropile (Jiggins 2017), implying selective pressures for increased visual function. *Heliconius* vision has been investigated using a variety of broad and narrow-band stimuli (Crane 1955, colored paper flowers; Swihart 1967, narrow band interference filters; Swihart 1972, narrow-spectrum color fibers; Zaccardi et al. 2006, narrow band interference filters). Available evidence demonstrates that *Heliconius* have true color vision in the long wavelength range (Zaccardi et al. 2006) but so far investigations in the short wavelength range have been limited.

Here we test whether *Heliconius erato* are capable of discriminating between narrow band wavelengths within in the UV range in the context of foraging. We use male and female *H. erato* butterflies, and as controls, male and female *H. melpomene* and *Eueides isabella* butterflies. Both *H. melpomene* and *E. isabella* lack a second UV opsin protein expressed in the eye but for different reasons: protein expression of UVRh2 was lost in *H. melpomene* (McCulloch et al. 2017), and *E. isabella* —a closely-related outgroup —never evolved a second UV opsin (Yuan et al. 2010). By confirming UV color discrimination in *H. erato* butterflies, and ruling it out in *H. melpomene* and *E. isabella*, we demonstrate the functional significance of their UV opsin duplication.

## METHODS

### Animals

Butterflies were purchased as pupae from the Costa Rica Entomological Supply (La Guácima, Costa Rica). The pupae were kept in a humidified chamber until they eclosed, then they were sexed and marked with a unique number. The butterflies were fed using a 10% honey solution with one bee pollen granule dissolved per 2 ml of solution. Butterflies were only allowed to feed on the positive stimulus during the training and testing. A total of 362 butterflies were used in the study, of which 200 were successfully trained and used in complete trials: 80 *H. erato* (40 females, 40 males); 80 *H. melpomene* (40 females, 40 males); and 40 *E. isabella* (20 females, 20 males).

### Behavioral experiments and apparatus

The experiments and training took place indoors in a mesh enclosure constructed from PVC pipes, measuring 1 m × 75 cm × 75 cm, and the room temperature was 24° C. The top of the enclosure was lined with 8 fluorescent tubes (Philips TLD 965 18 W; Eindhoven, The Netherlands). Our apparatus for training and experiments was based on a design described in Zaccardi et al. (2006) and previously used to test color vision in the monarch butterfly (Blackiston et al. 2011; see also Swihart and Swihart 1970, Weiss and Papaj 2003, Takeuchi et al. 2006, Rodrigues et al. 2010, Kinoshita and Arikawa 2014, and Drewniak et al. 2020 for other apparatus’ used in butterfly visual learning). It consists of two 3.0 cm diameter stimuli presented side-by-side, separated by 6 cm on two black platforms set on a larger black plate, measuring a total of 20 cm × 10 cm (see Figure 2 and Supporting Video 1). The apparatus was positioned vertically at the far end of the enclosure. Two wavelength stimuli were presented to the butterflies at a time. Light was emitted from two KL2500 Schott cold light sources (Mainx, Germany) into light guides held stable with a light guide holder. The light from each guide passed through a diffusor, a 10 nm narrow band-pass filter (Edmund Optics; Barrington, NJ, USA), and then through a transparent Plexiglass feeder disk (see Figure 3 in Zaccardi et al. 2006 for a diagram). For our experiments we used four narrow band-pass filters in paired choice tests: 380 nm versus 390 nm, and 400 nm versus 436 nm. We use 380 and 390 nm as the UV stimuli because the sensitivity curves of the two UV photoreceptors overlap in this range (McCulloch et al. 2016a) (Figure 1). If the butterflies have UV color discrimination using the UV and the violet photoreceptors together then we would expect that they would be able to discriminate between these two wavelengths. We also chose 400 nm and 436 nm as a control for color vision in all three species using the UV and blue photoreceptors. The light intensities for each wavelength were adjusted so that between these four wavelengths of light, the intensities for the experiments ranged from 9.56 × 10^15^ to 1.71 × 10^17^; quanta s^-1^ steradian^-1^ cm^-2^.

**Figure 1:**
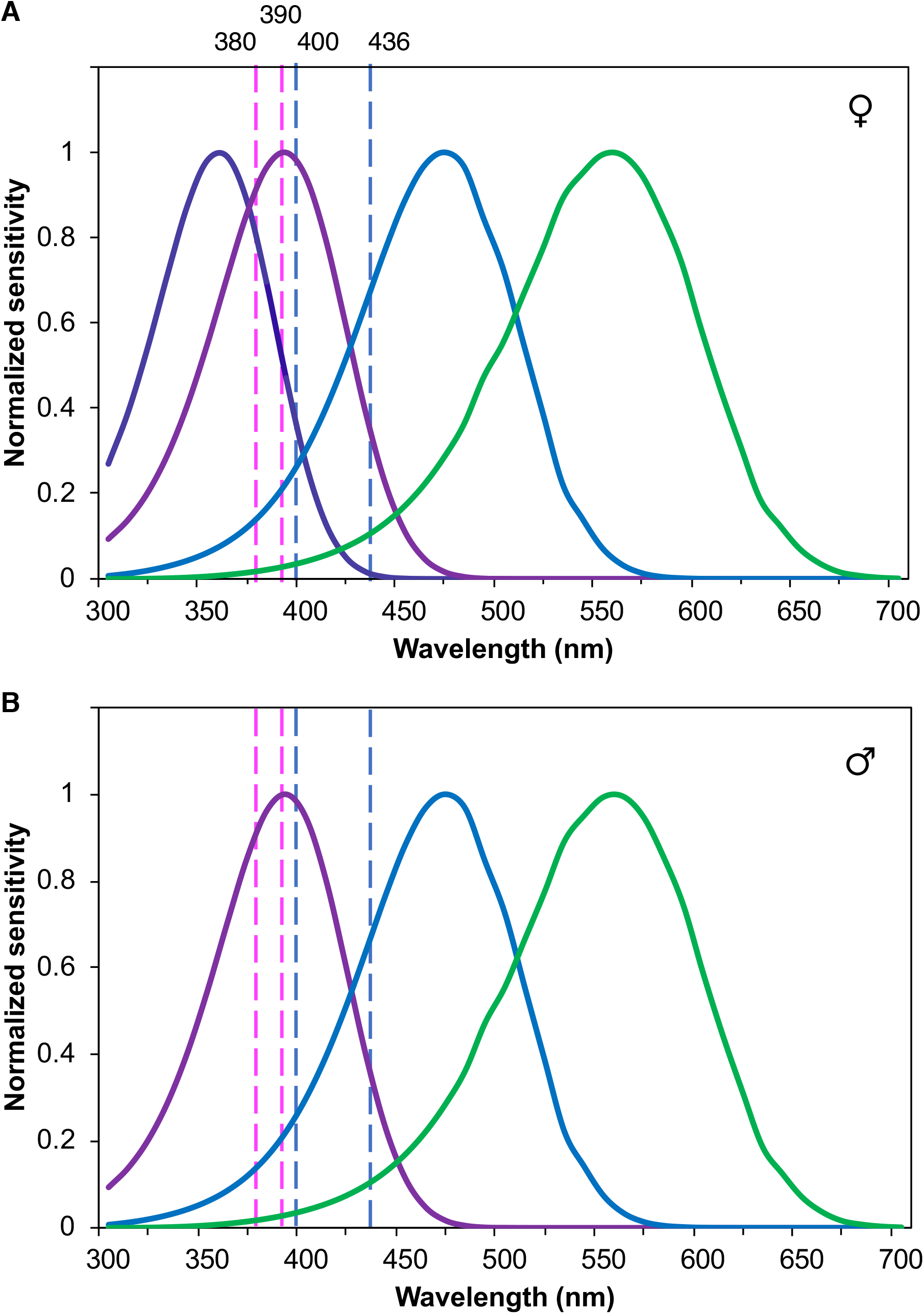
Normalized spectral sensitivities of the photoreceptors in adult (A) female and (B) male *H. erato* based on recorded intracellular spectral sensitivity maxima (McCulloch et al. 2016 *a*, *b*). The UV photoreceptor (dark purple), encoded by UVRh1, has a peak sensitivity at 355 nm, the violet photoreceptor (light purple), encoded by UVRh2, has a peak sensitivity at 390 nm, the blue photoreceptor (blue), encoded by the blue opsin has a peak sensitivity of 470 nm and the green photoreceptor (green), encoded by the LWRh opsin, has a peak sensitivity at 555 nm. A fifth known receptor class, with a peak at 600 nm due to filtering of the green rhodopsin by a red filtering pigment is not shown. Dotted lines represent the wavelength of peak transmission of the narrow bandpass fibers, 380 nm, 390 nm, 400 nm and 436 nm, used in discrimination tests. Male *H. erato* (B), lacking the UV photoreceptor (dark purple) are unable to discriminate between 380 and 390 nm light. *H. melpomene* and *Eueides isabella*, for different reasons, do not express the UVRh2 (light purple) opsin protein in their eye.

**Figure 2:**
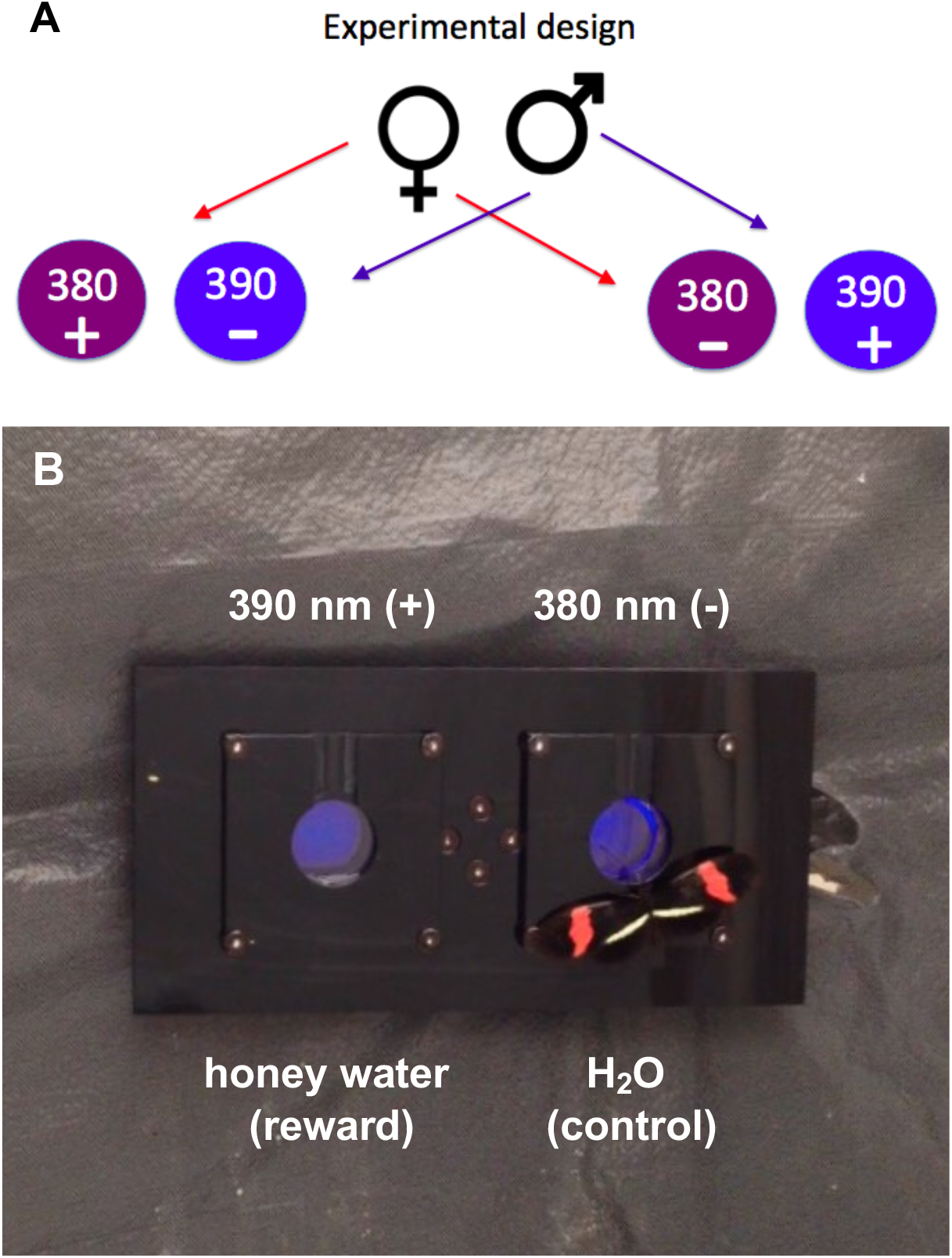
Experimental design of behavioral trials and experimental apparatus. (A) Female and male butterflies of three species, *Heliconius erato, H. melpomene* and *E. isabella* were reciprocally trained to associate honey water with a rewarded light (+) and tested using an apparatus (B) consisting of a rewarded light and an unrewarded light (-). Butterflies were trained and tested on their ability to discriminate 380 nm (right) from 390 nm (left) and 400 nm from 436nm lights (not shown). Shown is a male *H. erato* butterfly that has just landed on the light source apparatus during a trial.

**Figure 3:**
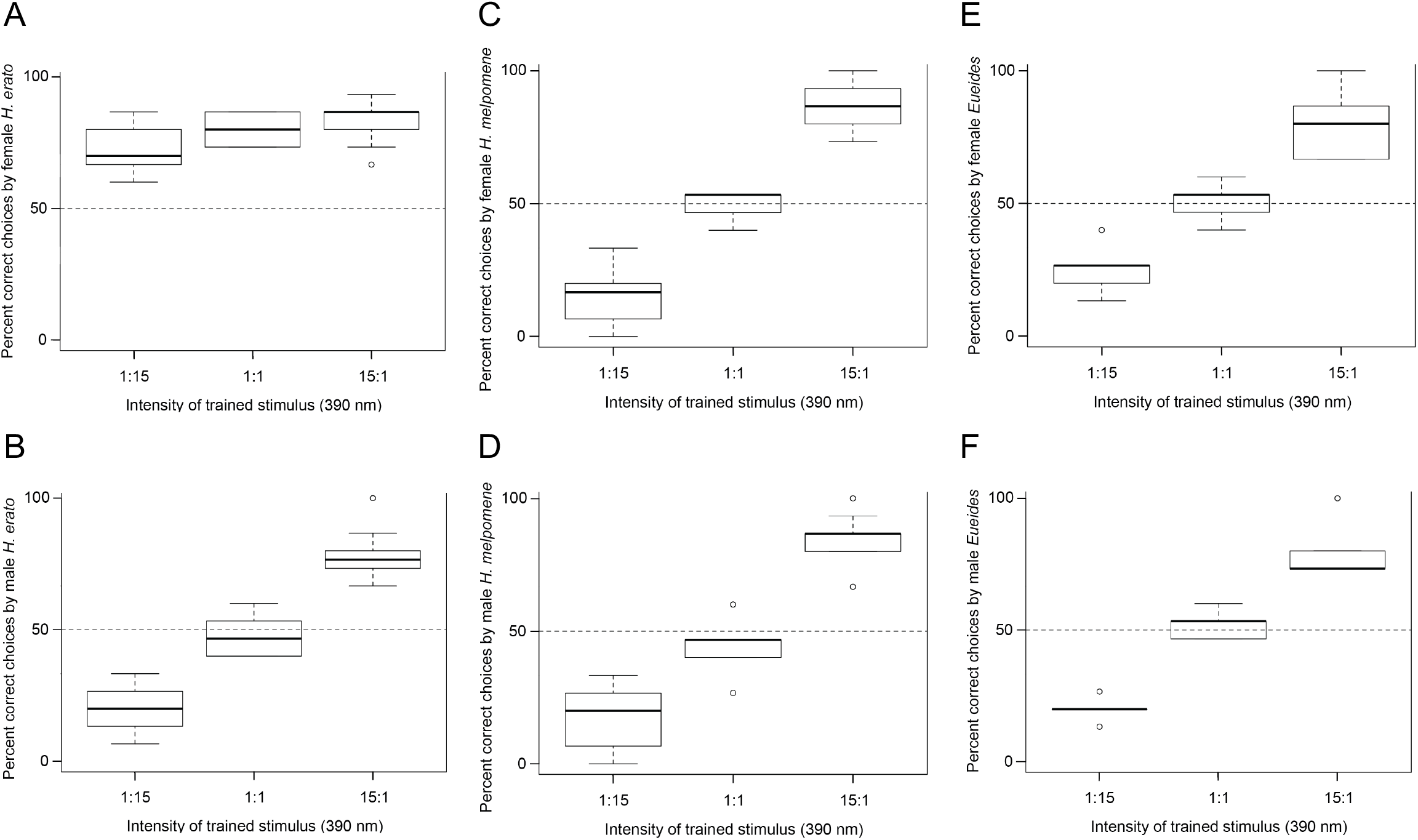
Percent correct choices to the rewarded (+) wavelength of 390 nm by *H. erato* females (A) and males (B), *H. melpomene* females (C) and males (D), and *E. isabella* females (E) and males (F) when given a choice between 390 nm (+) and 380 nm (-) light under varying intensities.

### Butterfly training

Butterflies were trained and fed for the first time within 15 hours of eclosion. Before training, they were allowed to acclimate to the experimental cage up to one minute, and only one butterfly was trained at a time. A droplet of food was placed in a small trough attached to the front of the feeder disk for the rewarded stimulus (+). The unrewarded stimulus (-) feeder trough remained empty. Each butterfly was trained by having its wings held together with forceps, and then slowly moved from the rear of the enclosure toward the apparatus to simulate a flying motion. The butterfly was then slowly waved in front of both the rewarded and unrewarded stimuli, and finally held in front of the rewarded stimulus where its proboscis was uncoiled with an insect pin until it came into contact with the food solution. At this point the butterfly would begin to drink. After the proboscis was manually uncoiled 2-3 times, the butterfly was able to uncoil the proboscis on its own in response to the stimulus. The procedure of carrying the butterfly with forceps from the rear of the cage to the light sources to feed was repeated 5 times per training session, with two training sessions per day separated by approximately 6 hours. Each time the butterfly fed from the rewarded stimulus, it was allowed to drink for 10 seconds, except for the very last training segment of the day where it was allowed to drink for several minutes. During training and between training sessions, the placement of the rewarded and unrewarded stimuli was randomly switched so that the butterfly did not learn to associate the left or right light with a food source. The apparatus was also cleaned thoroughly after each training session to minimize the association of chemical cues to the stimulus. After about 4-5 days of training, butterflies were capable of independently flying toward the apparatus when released from the rear of the cage and making a choice to fly to one of the two light stimuli (Supporting Video 1). At this point, the trained butterflies were starved for 20-24 hours then their choice trials began.

### Experimental trials

A separate cohort of butterflies was trained with each wavelength pair because the butterflies did not survive long enough to be trained multiple times. Both sexes of each species were first trained to 390 nm (+), and then tested for UV discrimination ability between 390 nm (+) and 380 nm (-) (ten per sex for *H. erato* and *H. melpomene* and five per sex for *E. isabella)*. The same number of individuals was trained to 380 nm (+) and given the choice between the two UV stimuli. Two new cohorts of butterflies were used for reciprocal training to 400 nm and to 436 nm. Three different approximate ratios of the physical intensities, or absolute brightnesses, of the +/- (rewarded/unrewarded) stimuli were used: 0.067, 1.0, and 15.0 (or 1:15, 1:1, and 15:1). The calculated ratios are 0.062, 1.0, 16.213 for 380 vs. 390 nm; and 0.0635, 1.0, 15.741 for 400 vs. 436 nm. These intensity ratios are described throughout the rest of this study as 1:15, 1:1, and 15:1, i.e. the rewarded stimulus (+) at 15 times less bright than the unrewarded stimulus (-), equal intensities for both stimuli, and the rewarded stimulus (+) at 15 times brighter than unrewarded stimulus (-). Butterflies first completed trials at an intensity combination of 1:1 (15 choices each). Following this test they were given random choices between intensities of 1:15 or 15:1 (rewarded:unrewarded) until they had completed 15 choices with each intensity combination.

The number of correct versus incorrect choices each butterfly made at different intensity combinations was modeled as dependent upon wavelength using general linear models in R statistical software (R Development Core Team 2010). We compared the ability of each category of butterfly to discriminate between the wavelength combinations at the different intensities. We also examined how discrimination abilities differed between all three butterfly species used in the study.

### Reflectance spectrometry

Live tissue was collected by accessing the butterfly and plant collection of Dr. Lawrence Gilbert at the Brackenridge Field Laboratory at the University of Texas, Austin on July 20, 2010. Reflectance spectra of *Heliconius erato petiverana* eggs*, Passiflora biflora* egg mimics, *Psychotria tomentosa* yellow infloresences, red bracts, and green leaves and *Psiguria warcewiczii* yellow and orange inflorescences and green leaves were measured by placing a probe holder (Ocean Optics RPH-1) over the specimen such that the axis of the illuminating and detecting fiber (Ocean Optics R400-7-UV/VIS) was at an elevation of 45 degrees to the plane of the tissue surface. Illumination was by a DH-2000 deuterium-halogen lamp, and reflectance spectra were measured with an Ocean Optics USB2000 spectrometer. Data were processed in MATLAB. Four to nine biological replicates per taxon were measured for each tissue type.

### Discriminability Modeling

To examine whether male or female *H. erato* eyes perform differently when viewing ecologically relevant objects, we constructed visual models. Models of color vision take into account how receptor signals contribute to chromatic (e.g., color opponent) mechanisms (Kelber et al. 2003). For *H. erato* males, we calculated discriminabilities for a trichromatic system consisting of UV2, blue and green receptors. For *H. erato* females, we calculated discriminabilities for a tetrachromatic system consisting of UV1, UV2, blue and green receptors. We excluded the red receptor from our calculations for both sexes because we do not have count data for this receptor class. Equations from Kelber et al. (2003) and Vorobyev & Osorio (1998) were used to model discriminabilities. This model incorporates a von Kries’s transformation, that is, normalization by the illumination spectrum, which models the way in which low-level mechanisms such as photoreceptor adaptation give color constancy (Kelber et al. 2003). Endler’s daylight illumination spectrum (Endler 1993) was used in the model. *H. erato* photoreceptor spectral sensitivity curves with *λ*_max_ values=355 nm (UV1)(female only), 390 nm (UV2), 470 nm (B), and 555 nm (L) from (McCulloch et al. 2016a) were used. Parameters for the butterfly visual models were as follows: Weber fraction=0.05 (Koshitaka et al. 2008) and relative abundances of photoreceptors, V=0.13, B=0.2, L=1 (male) or UV=0.09, V=0.07, B=0.17, L=1 (females) (McCulloch et al. 2016a).

## RESULTS

### UV discrimination

At the intensity of 1:1 for 390 and 380 nm light, female *H. erato* chose the rewarded light stimulus, 390 nm (+), significantly more than the unrewarded stimulus, 380 nm (-) (z-value = 6.791, p < 0.0001, Figure 3A). This indicates the ability of female *H. erato* to distinguish between the two UV wavelengths. The females continued to choose the correct, rewarded color stimulus under varying light intensity combinations. At an intensity ratio of 1:15 (rewarded: unrewarded), females significantly chose 390 nm (+) over 380 nm (-) (z-value = 5.19, p < 0.0001); and at an intensity of 15:1 (rewarded: unrewarded), females also chose 390 nm (+) over 380 nm (-) (z-value = 7.35, p < 0.0001). There was no difference between female preference for the correct stimulus with a 1:1 and 1:15 light ratio (z-value = −0.794, p = 0.427), or with a 1:1 and 15:1 light ratio (z-value = 0.319, p = 0.749), showing that females chose the correct light stimulus (390 nm) equally across all tested light intensity combinations.

With respect to male behavior, at the intensity of 1:1 for 390 (+) and 380 nm (-), male *H. erato* chose both the rewarded and unrewarded light stimuli equally (z-value = −0.49, p = 0.624, Figure 3B). This suggests they cannot distinguish between the two UV wavelengths. However, the males significantly preferred the correct, rewarded stimulus (390 nm)(+) when it was presented 15x brighter than the unrewarded stimulus (ratio 15:1 for rewarded: unrewarded; z-value = 6.421, p < 0.0001); and they significantly preferred the incorrect, unrewarded stimulus, 380 nm (-), at the intensity of 1:15 (rewarded: unrewarded; z-value = −6.671, p < 0.0001). These results imply that males prefer the brighter stimulus regardless of light wavelength, and further support their inability to discriminate between 390 and 380 nm. Comparing male and female performance, females significantly prefer the correct stimulus (390 nm)(+) more than males when 390 vs. 380 nm are at intensities of 1:1 (z-value = −3.427, p = 0.0006), and at intensities of 1:15 (z-value = −6.126, p < 0.0001), respectively. However, males and females equally chose the correct stimulus, 390 nm (+), when the rewarded:unrewarded intensity ratio was at 15:1 (z-value = −0.514, p = 0.607, Figure 3 A,B).

With *H. melpomene* and *E. isabella*, at the intensity of 1:1 for 390 and 380 nm, both sexes had similar wavelength discrimination behavior to male *H. erato* in that they chose both the rewarded (390 nm)(+) and unrewarded (380 nm)(-) light stimuli equally (z-value = 0.923, p = 0.356 for *H. melpomene*, Figure 3 C,D; z-value = 0.327, p = 0.744 for *E. isabella*, Figure 3 E,F). They were able to significantly choose the correct stimulus (390 nm)(+) only when it was 15x brighter than the unrewarded stimulus (z-value = −10.79, p < 0.0001 for *H. melpomene*; z-value = 6.791, p < 0.0001 for *E. isabella*), and they chose the unrewarded stimulus (380 nm)(-) significantly more when it was 15x brighter than the rewarded, correct stimulus (z-value = 10.460, p < 0.0001 for *H. melpomene*; z-value = −6.293, p < 0.0001 for *E. isabella*). No behavioral differences between sexes of either species were detected with statistical analyses (all p > 0.05), indicating that discrimination ability was consistent between both males and females of *H. melpomene* and *E. isabella*.

For the reciprocally rewarded tests, female *H. erato* butterflies were again consistent in discriminating between the rewarded (380 nm)(+) and unrewarded (390 nm)(-) stimuli when intensities were the same (z-value = −6.671, p < 0.0001, Supporting Figure 1A), when the rewarded stimulus was 15x brighter (z-value = −7.793, p < 0.0001), and when the rewarded stimulus was 15x less bright (z-value = −5.194, p < 0.0001). Male *H. erato* butterflies were incapable of discriminating between the different wavelengths when presented at equal intensities (z-value = −0.327, p = 0.744, Supporting Figure 1B), and chose the incorrect stimulus when it was 15x brighter than the correct, rewarded stimulus (z-value = 6.162, p < 0.0001). Males did, however, choose the correct stimulus when presented at an intensity ratio of 15x brighter than the unrewarded stimulus (z-value = −5.194, p < 0.0001). Females correctly chose the rewarded stimulus (380 nm)(+) significantly more than males at intensity ratios of 1:1 (z-value = −2.976, p = 0.00292) and 1:15 (z-value = −5.793, p < 0.0001), but at a ratio of 15:1 male and female *H. erato* chose the correct wavelength at similar rates (z-value = −1.424, p = 0.154, Supporting Figure 1 A,B).

Like male *H. erato*, *H. melpomene* and *E. isabella* could not distinguish between the two UV wavelengths presented at a 1:1 intensity ratio (z-value = 0.462, p = 0.644 for *H. melpomene*, Supporting Figure 1 C,D; z-value = 0.327, p = 0.744 for *E. isabella*, Supporting Figure 1 E,F). They significantly preferred the rewarded stimulus only when 15x brighter (z-value = −11.12, p < 0.0001 for *H. melpomene*; z-value = −7.024, p < 0.0001 for *E. isabella*), and preferred the unrewarded stimulus also only when 15x brighter (z-value = 7.793, p < 0.0001 for *H. melpomene; z*-value = 7.346, p < 0.0001 for *E. isabella*). Male and female discrimination behavior did not differ between *H. melpomene* or *E. isabella* (p > 0.05). In summary, female *H. erato* always discriminated between 380 and 390 nm light, consistently preferring the correct, rewarded stimulus, whereas male *H. erato*, male and female *H. melpomene*, and male and female *E. isabella* struggled with UV discrimination and only chose the correct stimulus when it was at a brighter intensity than the incorrect, unrewarded stimulus.

### Short wavelength discrimination

To investigate color vision in the blue range, we repeated the series of discrimination tests using 400 nm and 436 nm which would allow short wavelength discrimination using a UV or violet photoreceptor and a blue photoreceptor. As expected, when trained to 400 nm (+), female *H. erato* chose the correct stimulus when offered both light wavelengths at equal intensities (z-value = −7.93, p < 0.0001, Figure 4A), at an intensity of 1:15 for rewarded:unrewarded (z-value = −7.54, p < 0.0001), and at an intensity of 15:1 of rewarded:unrewarded light (z-value = −8.099, p < 0.0001). Male *H. erato*, male and female *H. melpomene*, and *E. isabella* behavior paralleled female discrimination behavior between the two blue wavelengths, with male *H. erato* choosing the correct wavelength at intensity combinations of 1:1 (z-value = −7.93, p < 0.0001, Figure 4B), 1:15 (z-value = −7.54, p < 0.0001), and 15:1 (z-value = −7.987, p < 0.0001); and *H. melpomene* and *E. isabella* males and females also choosing the correct, rewarded wavelengths at intensity ratios of 1:1 (z-value = −11.46, p < 0.0001 for *H. melpomene*, Figure 4 C,D; z-value = −7.63, p < 0.0001 for *E. isabella*, Figure 4 E,F), 1:15 (z-value = −11.07, p < 0.0001 for *H. melpomene*; z-value = −6.671, p < 0.0001 for *E. isabella*), and 15:1 (z-value = −11.47, p < 0.0001 for *H. melpomene;* z-value = −7.445, p < 0.0001 for *E. isabella*).

**Figure 4:**
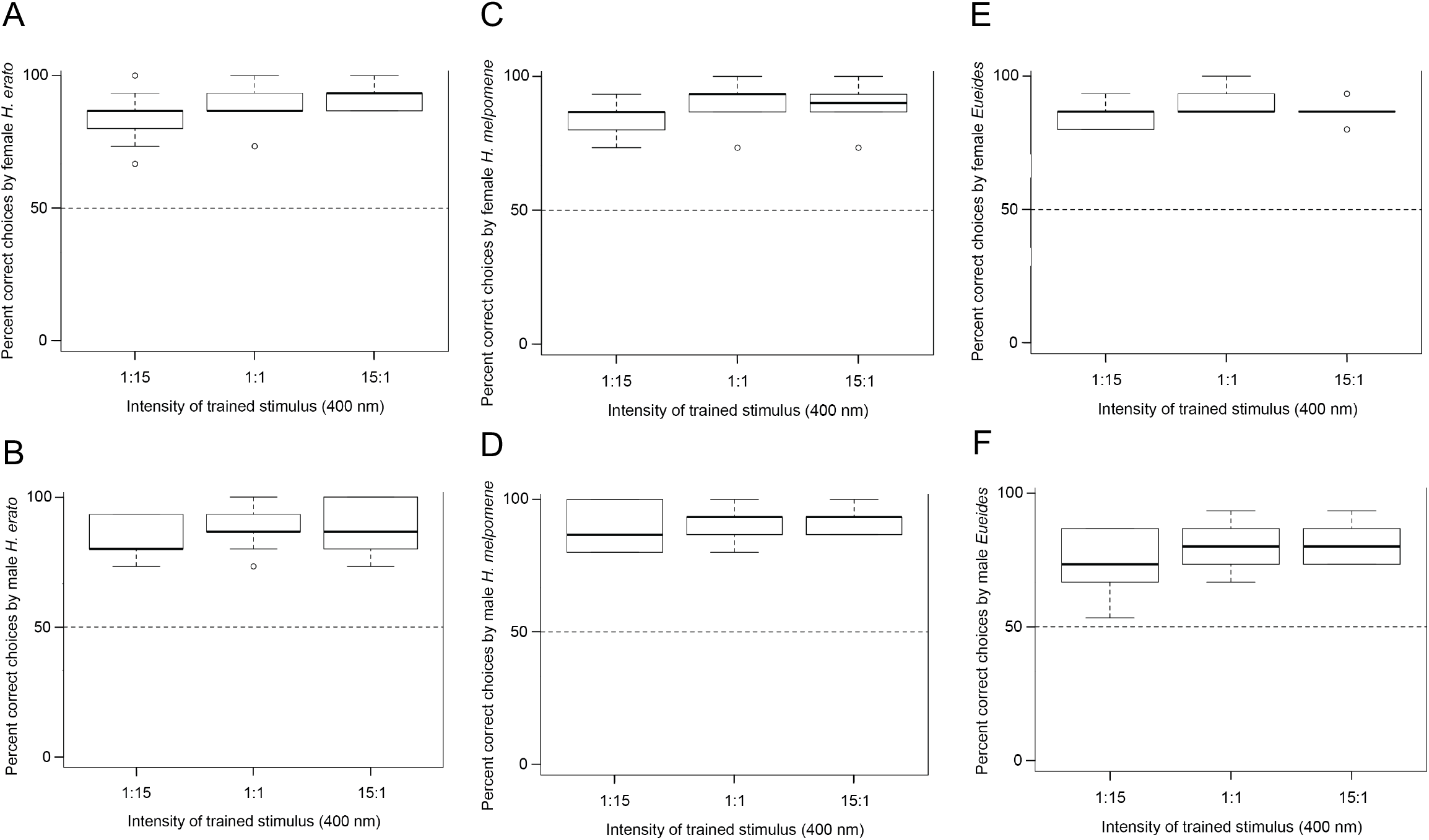
Percent correct choices to the rewarded (+) wavelength of 400 nm by *H. erato* females (A) and males (B), *H. melpomene* females (C) and males (D), and *E. isabella* females (E) and males (F) when given a choice between 400 nm (+) and 436 nm (-) light under varying intensities.

When trained to 436 nm (+), all butterflies continued to show a significant preference for the correct wavelength stimulus regardless of intensity. Female and male *H. erato* preferred the rewarded stimulus at equal intensities (z-value = 7.930, p < 0.0001 for females, Supporting Figure 2A; z-value = 7.714, p < 0.0001 for males, Supporting Figure 2B), at an intensity combination of 1:15 (z-value = 7.242, p < 0.0001 for females; z-value = 6.909, p < 0.0001 for males), and at 15:1 (z-value = 7.987, p < 0.0001 for females; z-value = 7.865, p < 0.0001 for males). *H. melpomene* and *E. isabella* followed the same trend and significantly preferred the correct wavelength (436 nm) (+) at an intensity combination of 1:1 (z-value = −10.85, p < 0.0001 for *H. melpomene* Supporting Figure 2 C,D; z-value = 7.793, p < 0.0001 for *E. isabella*, Supporting Figure 2 E,F), 1:15 (z-value = −9.853, p < 0.0001 for *H. melpomene*; z-value = 6.293, p < 0.0001 for *E. isabella*), and 15:1 (z-value = −11.07, p < 0.0001 for *H. melpomene*; z = 7.930, p < 0.0001 for *E. isabella*). There was no difference between *H. erato* male and female behavior, between *H. melpomene* male and female behavior, or between *H. erato, H. melpomene*, and *E. isabella* behavior (all p > 0.05) for selecting the correct light wavelength when trained to either 400 nm or 436 nm. All butterflies expressed the same ability to discriminate between 400 nm and 436 nm across all three intensity combinations.

## DISCUSSION

We conclude that *Heliconius erato* butterflies have true color vision in the UV range, between 380 nm and 390 nm, and that this is a female-limited behavior. Our results provide behavioral evidence that these butterflies can discriminate between more than one UV color using an ultraviolet and a violet photoreceptor, which suggests that the *UVRh1* (ultraviolet) and *UVRh2* (violet) opsin genes in *H. erato* function in the context of UV color discrimination. We also show that *H. erato, H. melpomene*, and *E. isabella* have color vision in the blue range between 400 nm and 436 nm, using both an UV and blue photoreceptor.

True UV color discrimination in *H. erato* is possible because of the evolution of a violet-sensitive photoreceptor, UVRh2, which has been present since the genus originated (Briscoe et al. 2010). As noted above, some clades (e.g., *H. melpomene)* do not express the UVRh2 protein at detectable levels in the adult compound eye despite expressing the *UVRh2* mRNA, due to ongoing pseudogenization (McCulloch et al. 2017). Opsin duplication events are not uncommon in butterflies (Sison-Mangus et al. 2006, Lienard et al. 2020). For example, the lycaenid butterfly *Polyommatus icarus* uses its duplicated blue opsin to see green, perhaps for discrimination of oviposition sites (Sison-Mangus et al. 2008). The pierid butterfly *Pieris rapae* has both a duplicated blue opsin and spectrally tuned filtering pigments: photoreceptor modifications that may be crucial for mate recognition by males (Arikawa et al. 2005; Wakakuwa et al. 2010). Yet another study has found that while both sexes of the wood tiger moth, *Arcia plantaginis* can distinguish between white and yellow male morphs (and females prefer to mate with white males), variation in female orange and red coloration is indiscriminable by both sexes, suggesting the moths’ visual system has evolved to facilitate female choice (Henze et al. 2018).

In *Heliconius erato* females, duplicate UV opsin genes encoding a UV and a violet receptor allow for UV color discrimination. The diversity of duplicated UV opsin presence or absence and spatial expression across the genus *Heliconius* is nonetheless thought-provoking. Male *H. erato* butterflies evidently use their duplicated UVRh2 (violet), blue, and long wavelength opsins in the context of mate choice discrimination of 3-OHK versus non-3-OHK yellow wing colors (Finkbeiner et al. 2017), an advantage predicted by modeling the discrimination abilities of *H. erato* males in comparison with a hypothetical male *H. erato* visual system in which UVRh1 takes the place of UVRh2 (Table 1) (McCulloch et al. 2017). Moreover, the loss of UVRh2 protein expression in *H. melpomene* (which use their ancestral UVRh1 opsin and not UVRh2) may contribute to increased attempts to mate with other species due to a reduction in visual ability to recognize conspecifics (Bybee et al. 2012, Dell’Aglio et al. 2019). *Heliconius* are part of a large mimicry complex that includes both unpalatable within-genus Müllerian mimics (which display 3-OHK yellow wing pigments) and palatable Batesian mimics such as *Eueides isabella* (which display unknown yellow wing pigments) (Srygley and Chai 1990; Bybee et al. 2012). Consequently, *Heliconius erato* (but not *H. melpomene*) butterflies benefit from having the violet receptor, UVRh2, which facilitates discrimination of yellow pigments of mimics from those of conspecifics.

**TABLE 1.**
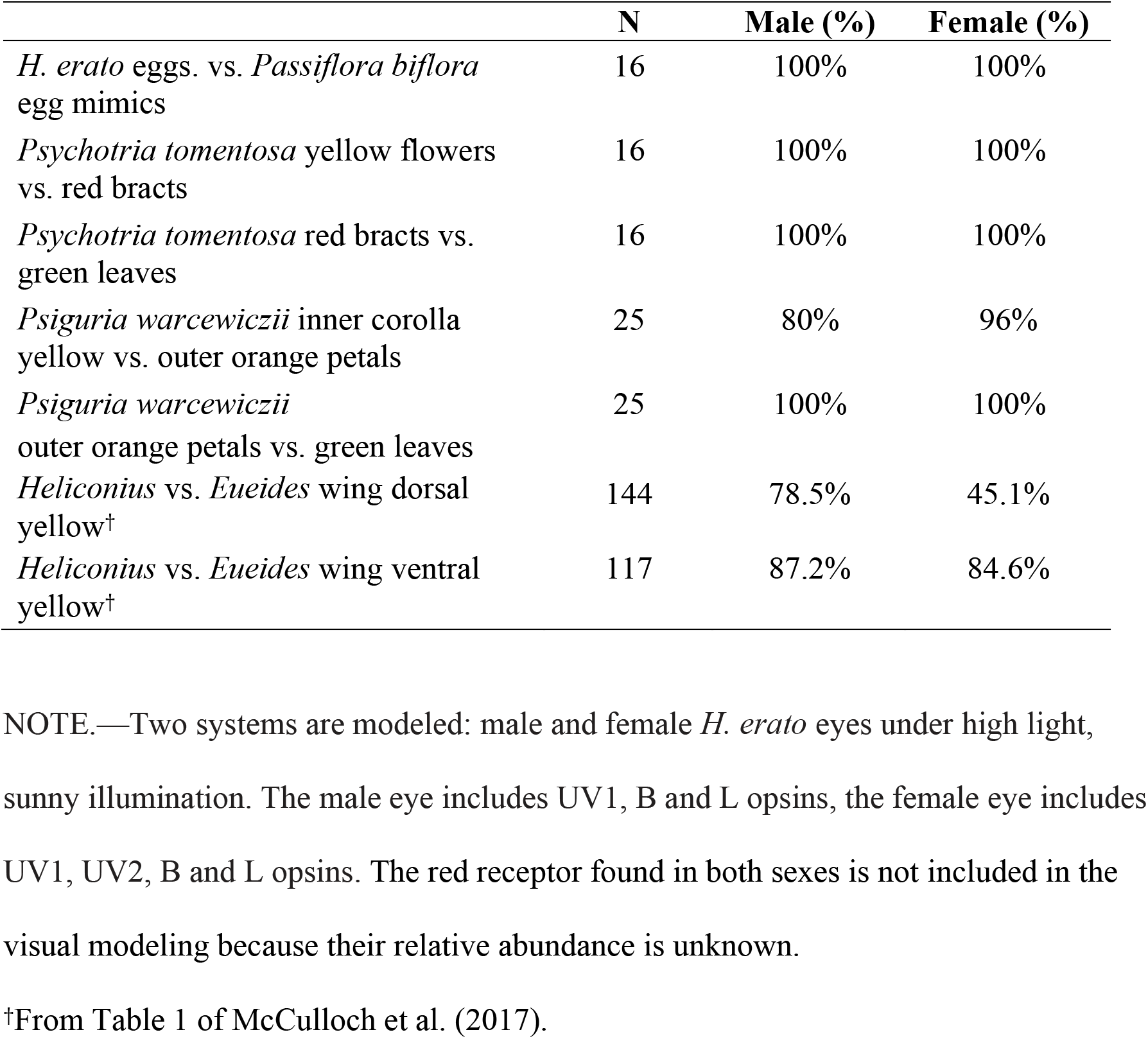
Percentage of *Heliconius* egg, egg mimic, pollen flower and wing colors with chromatic just noticeable difference (JND) values >1 for male and female *H. erato* eyes.

Early visual modeling of the *Heliconius* visual system suggested an additional benefit to *Heliconius’* displaying 3-OHK yellow pigments on the wing: with a second UV opsin in their eyes, more colors can be discriminated among *Heliconius* yellows than can be discriminated among the yellows of outgroup taxa (Briscoe et al. 2010). More recent work suggests *Heliconius* species may indeed be more conspicuous to conspecifics in their preferred habitats and light environment (Dell’Aglio et al. 2018, Dell’Aglio et al. 2019).

Both *H. erato* and *H. melpomene* may interact together by forming communal roosts in the same home range, which would provide added anti-predatory benefits through a similar visual signal (Finkbeiner et al. 2012). *Heliconius* co-mimics have been observed foraging together (pers. obs.) and roosting together (although uncommon; Mallet 1986, Finkbeiner 2014), and this could represent one instance where identifying a *Heliconius* individual (whether or not a co-species) would be beneficial. Aside from visual signals, *Heliconius* frequently use pheromone cues for conspecific recognition, especially for short-range signaling, for example during courtship behavior (Estrada and Jiggins 2008, Darragh et al. 2017, van Schooten et al. 2020).

It is possible that the adaptive function of UV color discrimination in female *H. erato* butterflies extends beyond intra- and interspecific communication to include host plant or pollen plant recognition. Within *Heliconius*, different species are specialists on *Passiflora* host plants for oviposition, and some of these *Passiflora* species contain extrafloral nectaries that resemble yellow *Heliconius* eggs (Williams and Gilbert 1981). *Heliconius* are known to avoid ovipositing on host plants that already have eggs because larvae have cannibalistic tendencies (Brown 1982, De Nardin and Araújo 2011), and fresh, new shoots that are the most edible for larvae can be of limited quantity (Gilbert 1982). While it is possible that the egg mimic structures differ spectrally from actual eggs in their UV reflectance, thus potentially allowing the additional UV opsin to provide discrimination between natural and mimic eggs, our preliminary investigation of the reflectance spectra of *H. erato petiverana* eggs and *Passiflora biflora* egg mimics, indicates that there is little to no UV reflectance for either the eggs or the egg mimics (Figure 5, top). Moreover, visual models indicate that both male and female *H. erato* visual systems are both able to discriminate *H. erato* eggs from *Passiflora biflora* egg mimics (Table 1), and *P. biflora* egg mimics from *P. biflora* leaves but not *H. erato* eggs from *P. biflora* leaves (Figure 5, bottom).

**Figure 5:**
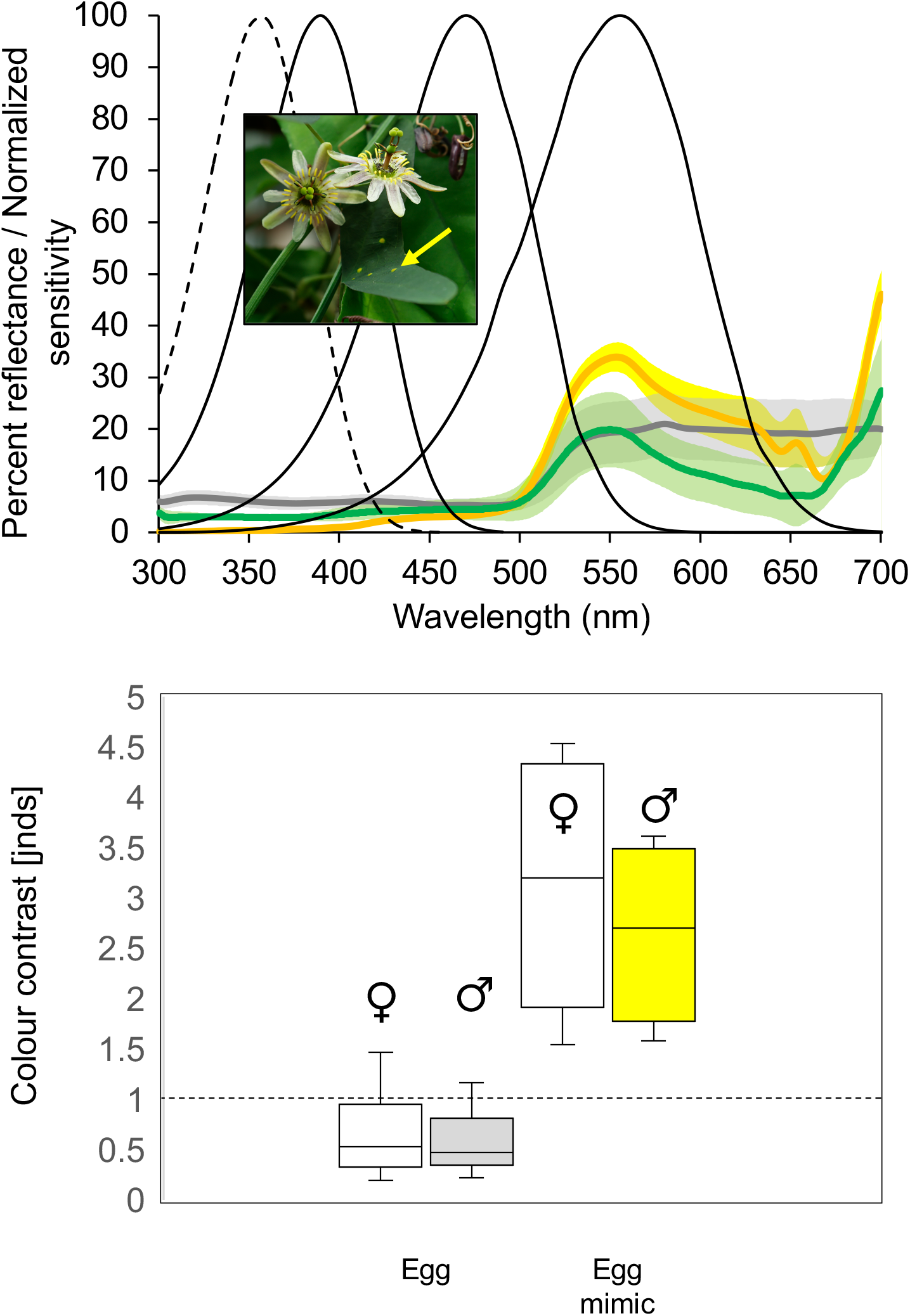
Reflectance spectra (top) and color contrasts (bottom) of *H. erato* eggs (dark grey line) and egg mimics (orange line) found on the leaves (green line) of *H. erato* host plant, *Passiflora biflora*. Shaded areas correspond to 95% confidence intervals, N=4. Black lines indicate UV1, UV2, blue, and green photoreceptors’ normalized spectral sensitivities. Not shown is the red receptor that is the result of filtering the green receptor with a red filtering pigment. Data from McCulloch et al. (2016 *a*, *b*). Bottom: Color contrasts between *H. erato* eggs and *P. biflora* leaves (N=16)(left) and between *P. biflora* egg mimics and *P. biflora* leaves (N=16)(right) in just noticeable differences (JNDs). Whiskers correspond to upper and lower limits. The absolute threshold is 1 JND; however in butterflies, the receptor noise levels are not well known so this is an approximation. Inset: *P. biflora* photograph with yellow arrow indicating egg mimic by C T Johansson. Source: Wikimedia: CC BY (https://creativecommons.org/licenses/by/3.0).

There is also the possibility that the leaves of caterpillar host plants, or even the petals of adult pollen flowers (such as *Psychotria* and *Psiguria)* have unique spectral properties in the UV range that would make a second UV/violet opsin beneficial. Intriguingly, we found evidence of a UV component to the reflectance spectra of the yellow inflorescences of *Psychotria tomentosa*, a plant from which *Heliconius* prolifically collect pollen (Figure 6, top). Both male and female *H. erato* visual systems appear adept, however, at discriminating between the yellow inflorescence from the red bracts of *Psychotria tomentosa* and at discriminating the red bracts from the green *Psychotria* leaf (Table 1, Figure 6, bottom). We also found that the yellow inner part of the *Psiguria warcewiczii* inflorescence has an even brighter UV component (Figure 7, top). Notably, the female *H. erato* visual system seems to have a bit of an advantage over the male *H. erato* visual system in discriminating the inner yellow from the outer orange petals of *Psiguria warcewiczii* flowers (Table 1, Figure 7 bottom). This difference is intriguing in light of evidence that female *Heliconius charitonia* (which have similar visual systems to *H. erato*)(McCulloch et al. 2017) collect significantly more pollen than do male *H. charitonia* because of their higher protein requirements for egg production (Boggs 1981; Boggs et al. 1981; Cardoso 2001; Estrada and Jiggins 2002; Mendoza-Cuenca and Macías-Ordóñez 2005); *H. charitonia* also display a sexual dimorphism in the flowers they collect pollen from with females preferring *Hamelia patens* pollen and males preferring *Lantana camara* flowers in one study locality (Mendoza-Cuenca and Macías-Ordóñez 2005).

**Figure 6:**
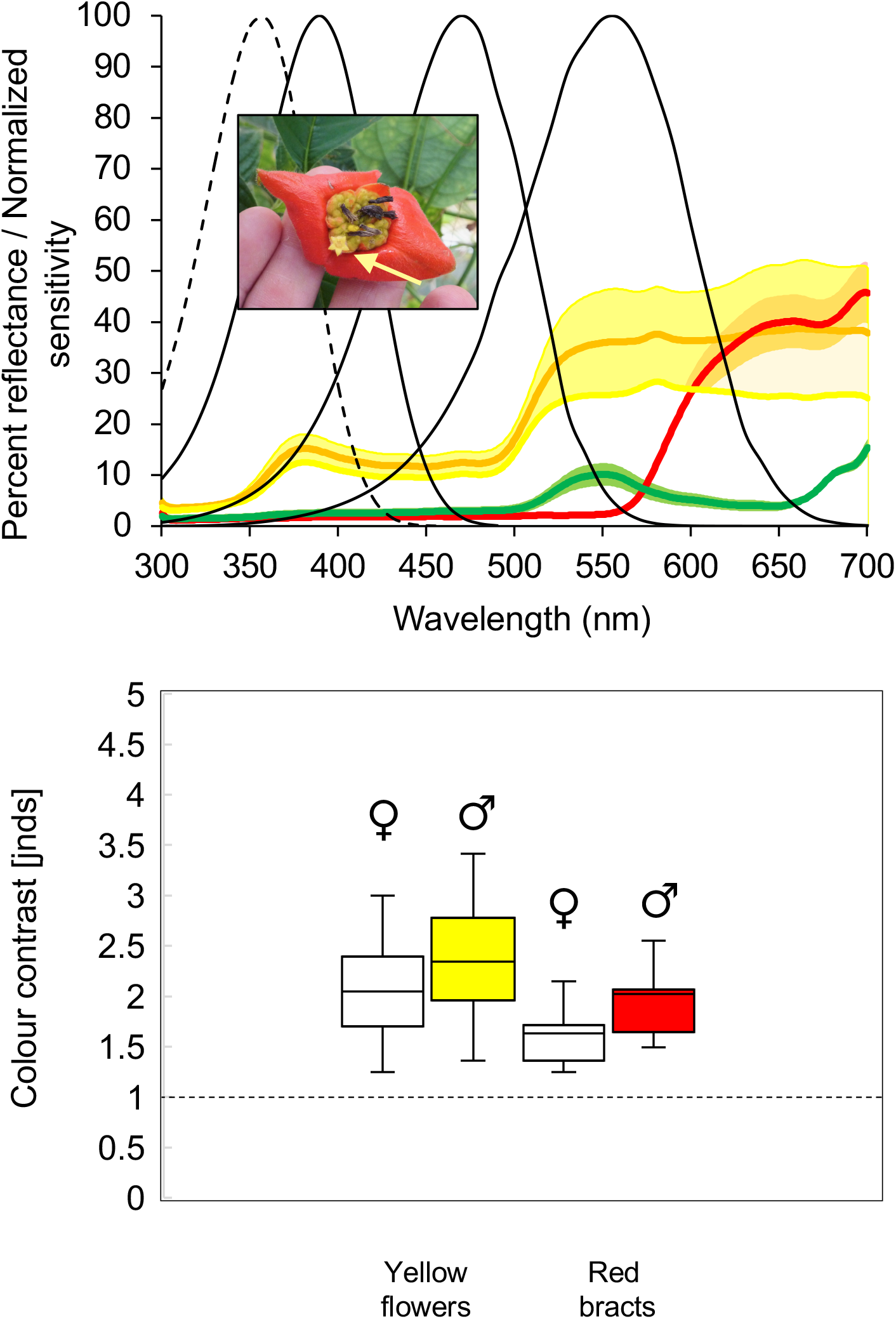
Reflectance spectra (top) and color contrasts (bottom) of *Psychotria tomentosa* yellow inflorescence (yellow line), red bracts (red line) and green leaves (green line), a plant from which *Heliconius* butterflies collect pollen. Shaded areas correspond to 95% confidence intervals, N=4-5. Black lines indicate UV1, UV2, blue, and green photoreceptors’ normalized spectral sensitivities. Data from McCulloch et al. (2016 *a*, *b*). Bottom: Color contrasts between yellow inflorescence and red bracts (N=16)(left) and red bracts and green leaves (N=16)(right) in just noticeable differences (JNDs). Whiskers correspond to upper and lower limits. The absolute threshold is 1 JND; however in butterflies, the receptor noise levels are not well known so this is an approximation. Inset: *P. tomentosa* photograph with yellow arrow indicating the yellow inflorescence. Surrounding the inflorescence are red bracts.

**Figure 7:**
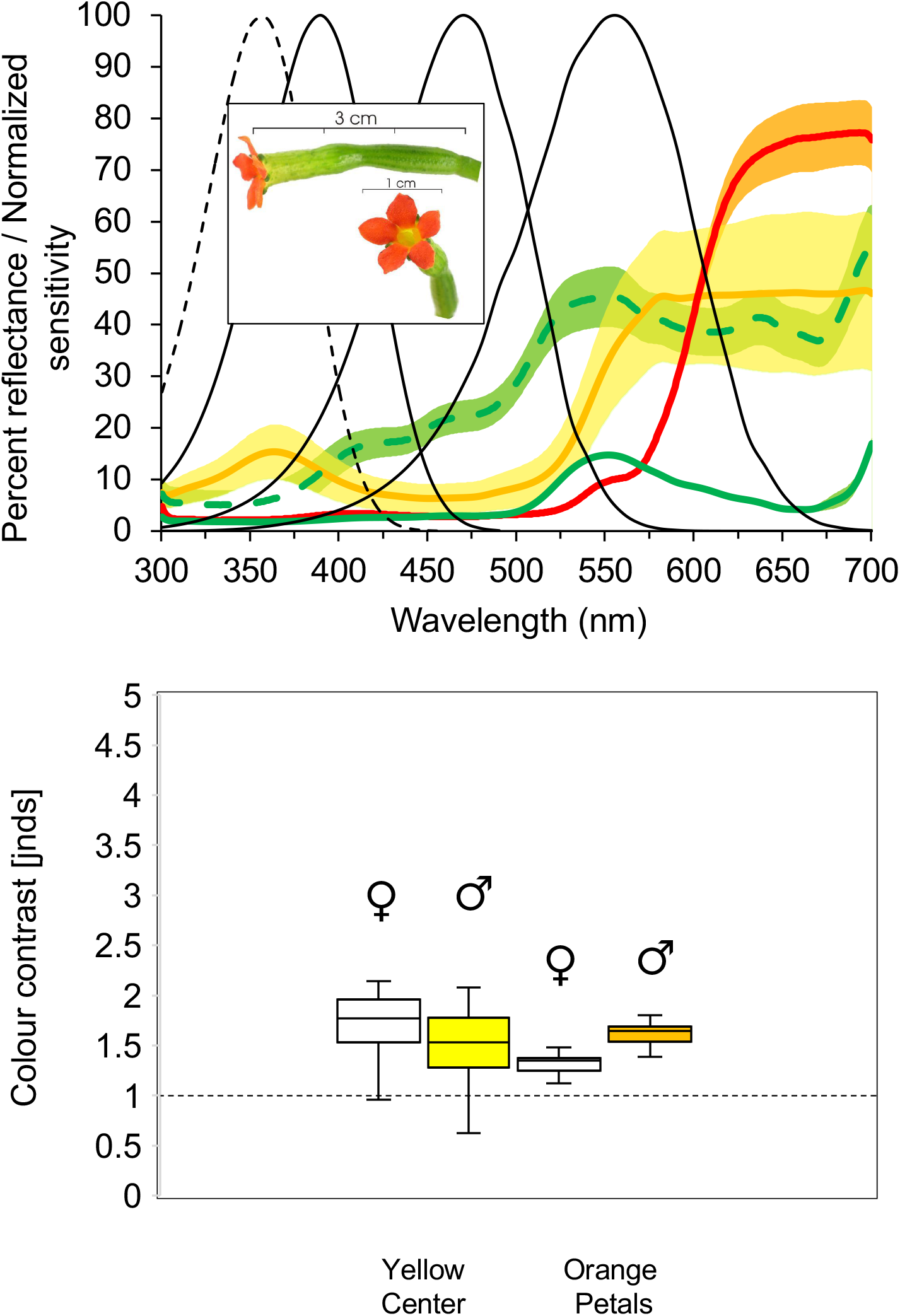
Reflectance spectra (top) and color contrasts (bottom) of *Psiguria warcewiczii—*yellow flower center (yellow line), outer orange petals (red line), light green corolla (dotted green line) and green leaves (solid green line)—a plant from which adult *Heliconius* butterflies collect pollen. Shaded areas correspond to 95% confidence intervals, N=5-9. Black lines indicate UV1, UV2, blue, and green photoreceptors’ normalized spectral sensitivities. Data from McCulloch et al. (2016 *a*, *b*). Bottom: Color contrasts between yellow flower center and outer orange petals (N=25) (left) and between outer orange petals and green leaves (N=25) (right) in just noticeable differences (JNDs). Whiskers correspond to upper and lower limits. The absolute threshold is 1 JND; however in butterflies, the receptor noise levels are not well known so this is an approximation. Inset: *P. warcewiczii* photograph. Photo credit: Steven Paton, Smithsonian Tropical Research Institute. Reprinted with permission.

An additional area ripe for exploration although not considered in the present study is in the investigation of ultraviolet polarized light cues in the context of host plant recognition. At least two studies of butterfly oviposition behavior have found that *Papilio* and *Pieris* butterflies respond to visible wavelength polarized light cues (Kelber et al. 2001; Blake et al. 2020), and previous work on *Heliconius cydno* finds they are able to use polarized light as a mating cue (Sweeney et al. 2003). Extending future investigations of *Heliconius erato* behavior to include UV polarized cues in the context of oviposition and mate choice seems likely to yield further insights into selective forces driving the evolution of this visual system’s sexual dimorphism.

Other animals that have photoreceptor spectral sensitivity in the UV range likely have true UV color discrimination, although to rule out brightness discrimination further experimentation is needed. Hummingbird hawkmoths (*Macroglossum stellatarum*) can discriminate between 365 nm and 380 nm, but it is unclear whether they are able to do so by means of true color vision or an achromatic cue (Kelber and Henique 1999). A different study showed that these moths are indeed able to discriminate between long wavelength stimuli under a range of intensities (Telles et al. 2016). In the case of the mantis shrimp and similar stomatopods, whose compound eyes possess the largest number of photoreceptor types known in any animal (including four UV-sensitive photoreceptors, Marshall and Oberwinkler 1999), there is little indication that their photoreceptors function with respect to true color vision at all (Thoen et al. 2014). However, our study provides clear evidence that despite differences in light intensity, *H. erato* female butterflies have the ability to discriminate between two UV wavelengths, lending support to the hypothesis that the new UV opsin gene in *Heliconius* functions in the context of UV color discrimination, and is one of the first to show that an animal can see multiple UV wavelengths using true color vision. In conclusion, our current and prior findings strongly suggest that both sexual selection and sex-specific natural selection have shaped the sexually-dimorphic visual system of *Heliconius erato*.

## ACKNOWLEDGEMENTS

We are grateful to Lawrence Gilbert for permission to measure reflectance spectra from host plants and flowers housed in the Brackenridge Field Laboratory; Robert Reed, Kailen Mooney, and Nancy Burley for advice and aid in project design; Paola Vargas and the Costa Rica Entomological Supply for providing live butterflies for experiments; Kyle McCulloch and Aide Macias-Muñoz for manuscript feedback and assistance with live butterfly care; and our funding sources: the U.S. Department of Education GAANN Fellowship, and the National Science Foundation (NSF) Graduate Research Fellowship under award no. DGE-0808392 to S.D.F. and NSF grant no. IOS-1257627 and IOS-1656260 to A.D.B.

## SUPPORTING INFORMATION

Supporting Figure 1: Percent correct choices to the rewarded (+) wavelength of 380 nm by *H. erato* females (A) and males (B), *H. melpomene* females (C) and males (D), and *E. isabella* females (E) and males (F) when given a choice between 380 nm (+) and 390 nm (-) light under varying intensities.

Supporting Figure 2: Percent correct choices to the rewarded (+) wavelength of 436 nm by *H. erato* females (A) and males (B), *H. melpomene* females (C) and males (D), and *E. isabella* females (E) and males (F) when given a choice between 436 nm (+) and 400 nm (-) light under varying intensities.

Supporting Video 1: A male *H. erato* butterfly is shown flying towards, and landing, on the light source apparatus during a trial. The light wavelengths presented are 390 nm (left) and 380 nm (right). The male chose 380 nm, the unrewarded stimulus, while it was presented at 15x brighter than the rewarded light stimulus of 390 nm.

