## Supplementary figures and images for "True UV color vision in a butterfly with two UV opsins"

### Supporting Figure 1

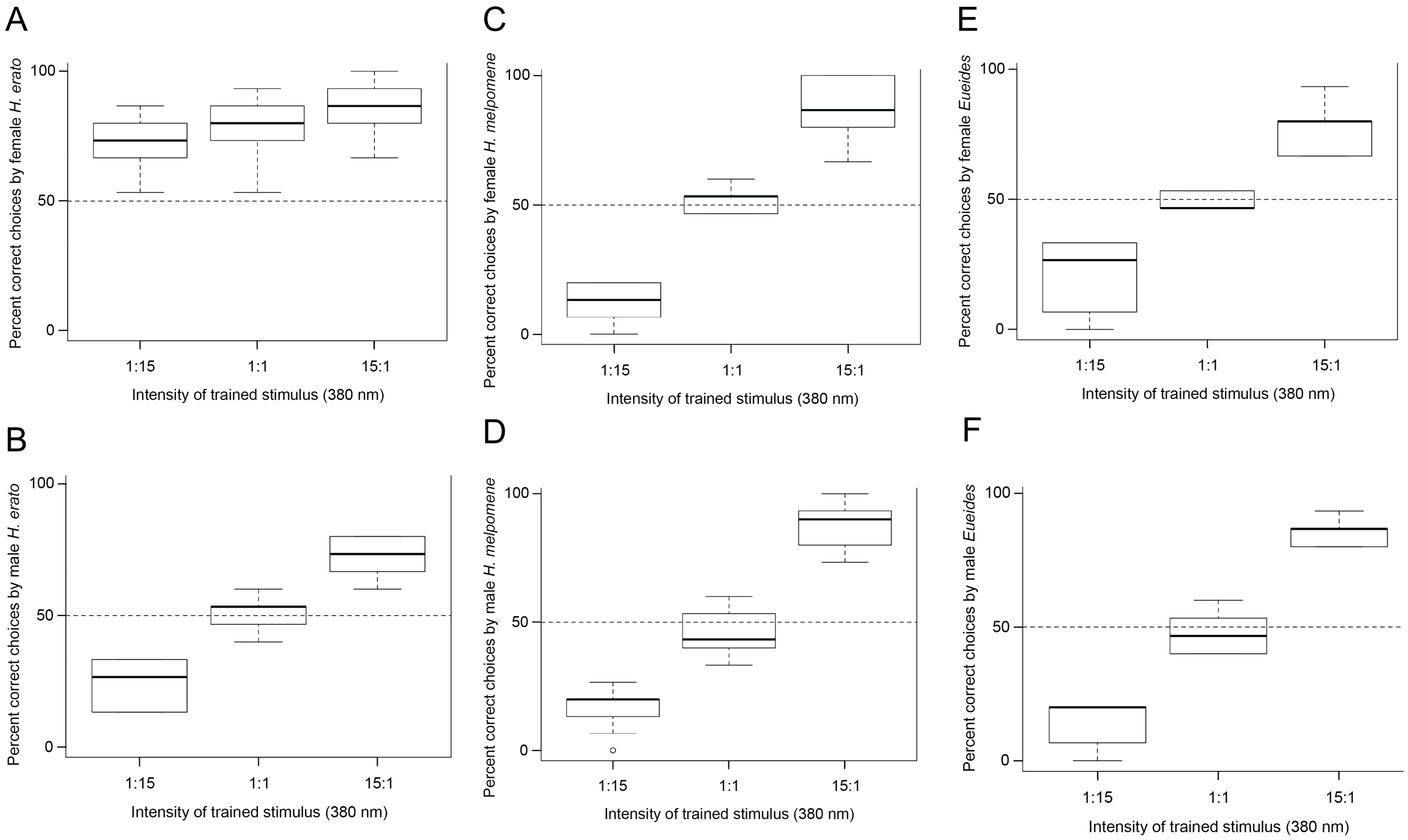


**Supporting Figure 1**

### Supporting Figure 2

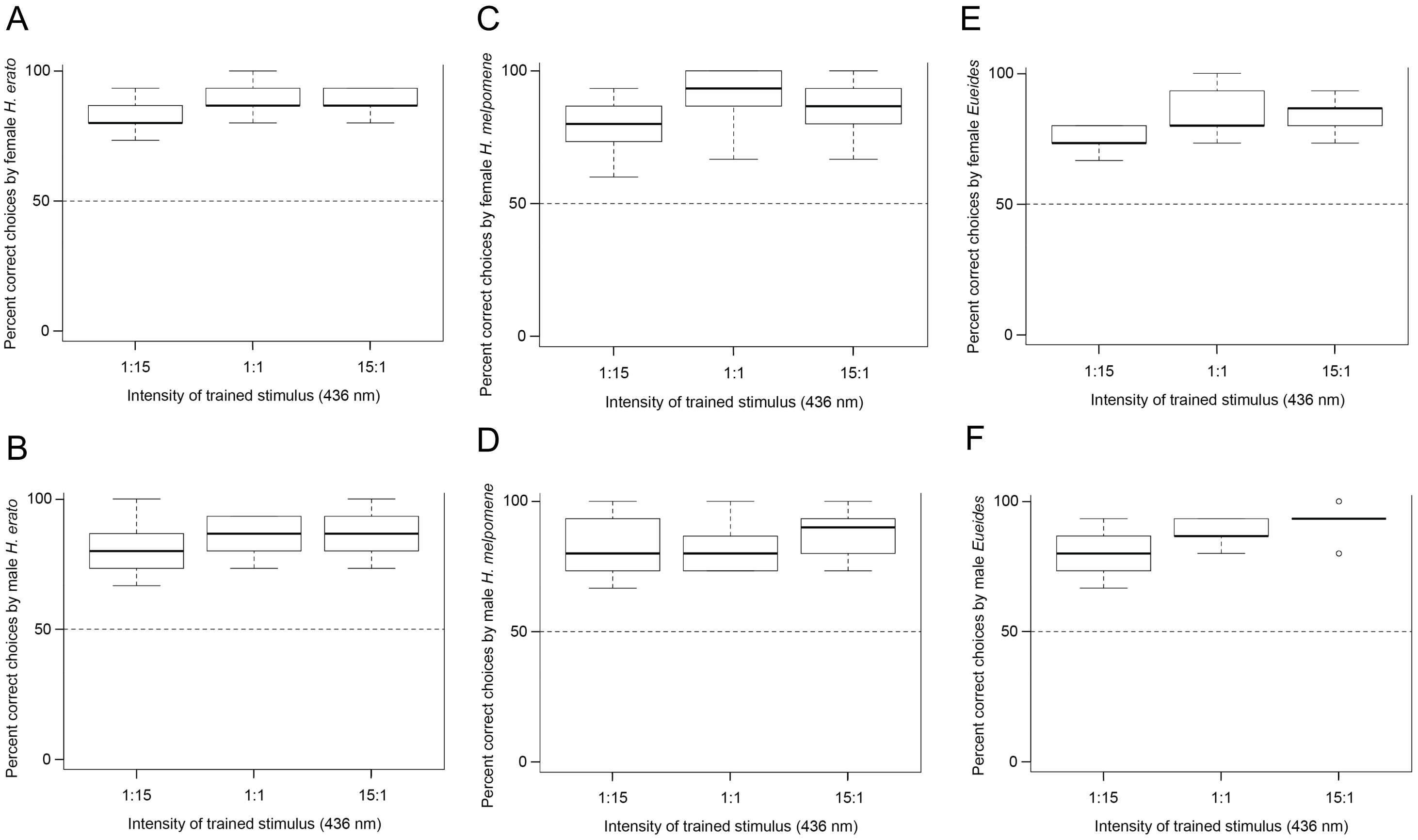


**Supporting Figure 2**
